# Dollo parsimony overestimates ancestral gene content reconstructions

**DOI:** 10.1101/2023.11.20.567878

**Authors:** Alex Gàlvez-Morante, Laurent Guéguen, Paschalis Natsidis, Maximilian J. Telford, Daniel J. Richter

**Affiliations:** Institut de Biologia Evolutiva (CSIC-Universitat Pompeu Fabra), Passeig Marítim de la Barceloneta 37-49, 08003 Barcelona, Spain; Université Claude Bernard Lyon 1, LBBE, UMR 5558, CNRS, Villeurbanne, 69622, France; Centre for Life’s Origins and Evolution, Department of Genetics, Evolution and Environment, University College London, Gower Street, London, WC1E 6BT

**Keywords:** ancestral reconstruction, Dollo parsimony, maximum likelihood, gene family evolution, phylogenomics

## Abstract

Ancestral reconstruction is a widely-used technique that has been applied to understand the history of gain and loss of gene families over evolutionary time scales, and to produce hypotheses on how these gains and losses may have influenced the evolutionary trajectories of extant organisms.

Ancestral gene content can be reconstructed via different phylogenetic methods, such as maximum likelihood or Bayesian inference, but many current and previous studies employ Dollo parsimony. We hypothesize that Dollo parsimony is not appropriate for ancestral gene content reconstruction inferences based on sequence homology, as Dollo parsimony is derived from the assumption that a complex character can only be gained once in evolutionary history. This premise does not accurately model molecular sequence evolution, in which false orthology can result from sequence convergence (including random sequence similarity and parallel gene gains), non-orthologous homology or lateral gene transfer. The aim of this study is to test Dollo parsimony’s suitability for ancestral gene content reconstruction and to compare its inferences with a maximum likelihood-based approach which allows a gene family to be gained more than once within a tree.

In order to test our hypothesis, we first compared the performance of the two approaches on a series of artificial datasets each of 5,000 genes that were simulated according to a spectrum of evolutionary rates. Next, we reconstructed protein domain evolution on a phylogeny representing known eukaryotic diversity. We observed that Dollo parsimony produced numerous ancestral gene content overestimations, which were more pronounced at nodes closer to the root of the tree. These observations led us to the conclusion that, confirming our hypothesis, Dollo parsimony is not an appropriate method for ancestral reconstruction studies based on sequence homology.

## INTRODUCTION

Ancestral reconstruction is the inference of ancient characteristics based on extant species characteristics across a phylogenetic tree relating those species. It can be applied to sequences (DNA, RNA or protein), as well as to morphological characters. It has been widely used to understand the history of gain and loss of gene families over long time scales, and to produce hypotheses on how these gains and losses may have influenced the evolutionary trajectories of extant organisms (Groussin et al., 2016). The power of this technique has allowed it to play a crucial role in diverse topics from tick evolution (Mans et al., 2016) to flower morphology and pollination (Pérez et al., 2006) to the unicellular-to-multicellular transition (Ros-Rocher et al., 2021) and many others (Harms & Thornton, 2010) (Kohn et al., 2006) (Sverdlov et al., 2004).

Ancestral reconstructions of gene family gain and loss can be based on different phylogenetic inference methods, such as maximum likelihood or Bayesian inference, but many current studies are based on Dollo parsimony.

Dollo parsimony is a specific case of maximum parsimony based on Dollo’s Law (Dollo, 1893), which was based on an interpretation of morphological characters and postulates that the same evolutionary path cannot be followed more than once, precluding the possibility that an identical character can be gained twice. This premise is implemented in Dollo parsimony by allowing characters to be gained only once, but accepting as many losses as necessary (Felsenstein, 1983).

The literature is replete with examples using Dollo parsimony as a phylogenetic inference method. One of the most frequent applications of Dollo parsimony has been in reconstructing the gains and losses of genes in the lineages leading to major multicellular eukaryotic groups, including land plants (Bowles et al., 2020), animals (Fairclough et al., 2013; Najle et al., 2023; Paps & Holland, 2018), and brown algae (Cock et al. 2010). It has been applied to examine patterns of gene gain and loss in the evolution of novel trophic modes or in the adaptation to specific environments in fungi and their relatives (Mikhailov et al. 2017, Galindo et al., 2018, Galindo et al. 2021, Mikhailov et al. 2022), in green algae (Repetti et al. 2022), and in red algae (Cho et al. 2023). Dollo parsimony has also been employed to investigate the evolution of gene gains and losses that may have led to physiological changes such as those underlying the evolution of *Wolffia australiana*, the smallest known flowering plant (Park et al., 2021).

In addition to analyses of gene gain and loss, Dollo parsimony has been applied to infer phylogenies in sweet cherry cultivars (Zhou et al., 2005), in *Mycobacterium* (Stevenson et al., 2002), and, using retroelements, in Laurasiatheria (a group of mammals) (Doronina et al., 2017). Dollo parsimony has also been used to reconstruct protein domain (Zmasek & Godzik, 2011) and intron (Csuros et al., 2011) gains and losses across the eukaryotic tree of life, and to study inverted repeat region structure, pseudogenization and gene loss in *Pedicularis*, a hemiparasitic land plant (in comparison with other reconstruction methods) (Li et al., 2021).

The basic assumption of a single gain of an orthologous gene family in Dollo parsimony is also implicit in phylostratigraphy, a widely used approach to reconstruct patterns of gene gain over evolutionary timescales, in which gene origins are assigned to the most recent common ancestor of the extant species in which the gene is found (Domazet-Loso et al., 2007; Domazet-Lošo & Tautz, 2010).

In ancestral gene content reconstruction studies, the standard process is first to use an orthology inference program such as OrthoFinder2 (Emms & Kelly, 2019), which use BLAST (or a BLAST-like software) to search for homology among the gene sequences of all input extant species (Altschul et al., 1990). Sequence similarity values are subsequently used as the basis to construct orthologous groups, generating an output (in the case of Dollo parsimony, a binary output representing the presence or absence of an orthologous group in a species) that is used as the input for the ancestral reconstruction programs.

Although Dollo parsimony is a practical method that can be appealingly simple when working with morphological characters, we hypothesize that it is not appropriate when the input data for ancestral reconstruction are derived from sequence homology. Dollo parsimony operates under the assumption that a feature can only be gained once. Under this assumption, if a gene is present in two different species anywhere in the analyzed phylogeny, it will always be inferred to have been present in their most recent common ancestor, even if the sequence similarity between the genes in the two species may have arisen by chance; the more the two species are distantly related, the more the origin of the gene will be pulled towards the root. This assumption does not take into account convergent sequence evolution (homoplasy) or horizontal gene transfer, and we posit that it results in an overestimation of gene losses and an underestimation of gene gains.

In order to test our hypothesis, we compared the ancestral gene content reconstructions produced by PHYLIP Dollop (Dollo parsimony) (Felsenstein, 1993) against Bppancestor (a maximum-likelihood method with a model of gene gain and loss in order to assess ancestral presence) (Guéguen et al., 2013) for a simulated dataset based on a phylogeny of metazoans. This dataset contained 200 independent simulations of the evolution of protein sequences over a fixed topology of 57 animal species (Natsidis et al., 2021). Each of these simulations contained 5,000 orthologous groups that were present in all 57 species, with no gains or losses allowed.

Next, we compared the reconstruction of Pfam protein domain evolution across the eukaryotic tree of life produced by Dollo parsimony versus maximum likelihood. We used the insights gained from our analysis of simulated data to compare the results of this reconstruction to a previous study, based on Dollo parsimony, which found that protein domain loss outweighed protein domain gain across eukaryotes, and that the last eukaryotic common ancestor possessed a protein domain repertoire larger than any extant species (Zmasek & Godzik, 2011).

## METHODS

### Simulated dataset: Input data

200 datasets derived from simulations of the evolution of protein sequences for a fixed topology of 57 species, representing metazoan phylogeny, were obtained from Natsidis *et al* (Natsidis et al., 2021). Each of these simulations contained 5,000 sets of orthologs present in all 57 species, and no gains or losses were allowed during the evolution of the protein sequences. These different simulations differed from each other in the overall evolutionary rate (branch length multiplier from 0.2x to 10x) and used simulation-specific alpha parameters to simulate genes with various degrees of site rate heterogeneity (alphas from 0.4 to 1.6).

OrthoFinder2, a platform for comparative genomics (Emms & Kelly, 2019), was run by Natsidis *et al*. separately on all 200 sets of 285,000 protein sequences (5,000 protein sequences per species). OrthoFinder’s output was converted with Perl scripts into a binary format suitable for Bppancestor and PHYLIP Dollop, in which each orthogroup in each species was scored as either present or absent. The version used was Perl 5.30.0 (Wall, L. et al., 2000).

### Simulated dataset: Running Bppancestor

Using Perl scripts we generated one configuration file per each input file, using a template (template_bppancestor_config_file.conf, available in the GitHub repository). The configuration files specified a stationary process with a birth/death model and a gamma distribution of rate variation among sites, which was stated to remain homogeneous across all branches of the topology. We performed tests with a non-stationary model, which generated equivalent results (data not shown).

We ran Bppancestor iteratively to perform the ancestral reconstructions on each simulation and processed the results with Perl scripts to parse their results. The number of presences at each node was counted by treating the estimated probabilities as expected values and summing them across all sites. The used versions were Bio++ version 3.0.0 (Guéguen et al., 2013) and Perl 5.30.0 (Wall, L. et al., 2000).

### Simulated dataset: Running PHYLIP Dollop

We ran PHYLIP Dollop separately for each simulation. We used Phylip Dollop’s default options, except for the option “Search for the best tree”, which we disabled because we provided a fixed tree topology as input, and the option “Print States at all nodes”, which we enabled. All operations were performed under PHYLIP version 3.697 (Felsenstein, 1983).

### Simulated dataset: Running Mesquite

The phylogenetic analysis was carried out using the “Trace All Characters” option in the “Tree:Analysis” tab of the “Tree Block” section of the program; with default settings and inputting an initial states file and a fixed phylogenetic tree. The initial states file was inputted in Nexus format, while the tree file was inputted in Newick format. All the operations were performed under Mesquite version 3.61 (Maddison, W. P. and D.R. Maddison, 2023).

### Pfam domain content in the earliest eukaryotes: Input data

A dataset containing 993 protein sets representing eukaryotic diversity was obtained from EukProt v3 (Richter et al., 2022). Then, we ran InterProScan 5.56-89.0 (Jones et al., 2014) to detect Pfam domains in the protein sequences. We converted the results into a binary format suitable for Bppancestor and PHYLIP Dollop, indicating the presence or absence of each Pfam domain in each species. The version used was Perl 5.30.0 (Wall, L. et al., 2000).

### Pfam domain content in the earliest eukaryotes: Tree topology

We ran Gappa 0.8.2 (Czech et al., 2020) on the taxonomy obtained from EukProt v3 (Richter et al., 2022) in order to generate our initial input tree. We used the AfterPhylo perl script (Zhu, 2014) to truncate the names of the tree to 10 characters and resolved it using the “multi2di” R function, from R version 3.6.3 (R Core Team, 2022), in order to generate the final version of the tree for PHYLIP Dollop. Multiple different versions of the randomly resolved tree were tested and generated equivalent results (data not shown).

### Pfam domain content in the earliest eukaryotes: Running Bppancestor

To estimate branch lengths and model parameters, we ran Bppml (Guéguen et al., 2013), with a configuration file (template_bppml_config_file.conf, available in the GitHub repository), specifying a stationary process with a binary model and a gamma distribution of rate variation among sites, which was stated to remain homogeneous across all branches of the topology.

We also generated a configuration file for Bppancestor (domains_configuration_file.conf, available in the GitHub repository) with the most likely model estimated by Bppml.

We ran Bppancestor with this configuration file and treated the output with a Perl script to parse the results and count the number of gene gains and losses. The script sums the probabilities of the genes present at each node, treating each individual probability as an expected value of gene presence. It then compares the probability of each gene’s presence relative to the parent node. If the probability of presence in the child is higher than probability in the parent, it is considered as a gain and added to the sum of gains leading to the node; if the probability is lower, then it is considered as a loss and added to the sum of losses leading to the node. The used versions were Bio++ version 3.0.0 (Guéguen et al., 2013) and Perl 5.30.0 (Wall, L. et al., 2000).

### Pfam domain content in the earliest eukaryotes: Running PHYLIP Dollop

We ran PHYLIP Dollop and treated the output with a Perl script to parse the results and count the number of gene gains and losses. The script sums the number of genes estimated to be present at each node. If there is a change in gene presence relative to the parent, it is recorded as either a gain or a loss. The phylogenetic analysis was carried out using PHYLIP Dollop’s default options, except for the option “Search for the best tree”, which we disabled because we provided a fixed tree topology as input, and the option “Print States at all nodes”, which we enabled. All operations were performed under PHYLIP version 3.697 (Felsenstein, 1983) and Perl 5.30.0 (Wall, L. et al., 2000).

### Graphics and figure design

We used Rstudio 2023.3.0.386 (RStudio Team, 2020) iTOL 6 (Letunic & Bork, 2007) and Inkscape 1.1.1 (Inkscape Project, 2020) to produce and modify figures and phylogenetic trees.

## RESULTS

### Dollo parsimony overestimates ancestral gene content in a simulated dataset

We tested the performance of Dollo parsimony on a dataset containing 200 simulations of protein sequence evolution on a fixed topology of 57 species (Natsidis et al., 2021). Each of these simulations contained exactly 5,000 orthologous genes present in all 57 species, with no gains or losses allowed. The 200 simulations differed from each other in their rates of substitution and in the variation of rates among sites within each gene. Separately for each individual simulation, OrthoFinder2 (Emms & Kelly, 2019) was run in order to partition simulated gene sequences into orthologous groups (Natsidis et al., 2021).

After ancestral reconstruction, any ancestral nodes inferred to have contained either more than 5,000 genes or fewer than 5,000 genes would represent incorrect estimates of the number of orthologous groups used as input to ancestral reconstruction, in the ancestral reconstruction inference method itself, or both.

We began by examining the contents of the orthogroups to be used as input to ancestral reconstruction. The expected result from a correct partitioning of simulated gene sequences into orthologous groups would be 5,000 orthologous groups, each containing exactly 57 sequences (one from each species). The orthogroups for most simulations contained, on average, sequences from fewer than 57 species (Supplementary Figure 1). As a consequence, there were more than the expected 5,000 orthogroups per simulation (Supplementary Figure 2), and each of those orthogroups contained only a subset of the 57 species. Most species did not have a gene present in many orthogroups due the artificial expansion of the number of orthogroups, the generation of singletons (which are not part of any orthogroup in OrthoFinder2’s output) and the grouping of multiple genes from the same species in the same orthogroup, which resulted in an underestimated input value (fewer than 5,000 orthogroups) for most input nodes in most simulations (Supplementary Figure 3B). Although simulations with slower rates of evolution were generally correctly partitioned into complete orthologous groups, as the simulated rate of evolution increased, so did the underestimation in the number of orthogroups present in each input species. This effect is likely the result of higher sequence divergence making it less likely that genes could be correctly grouped by homology into orthogroups. Since the simulated rates of evolution span the values likely to be present in real data sets (Natsidis et al., 2021), the composition of orthologous groups we used as input to ancestral reconstruction should be reflective of the scope of potential underestimates present in real data sets.

In order to determine whether Dollo parsimony performs similarly to other ancestral reconstruction methods that do not share its strict assumptions, we analyzed the same data set with a maximum likelihood method, Bppancestor (Guéguen et al., 2013). To exclude the possibility that our particular choice of maximum likelihood software might influence our results, we compared Bppancestor to another implementation, Mesquite (Maddison, W. P. and D.R. Maddison, 2023) on a subset of the input data. Bppancestor and Mesquite produced nearly identical ancestral reconstructions (Supplementary Figure 4). As Bppancestor can be easily automated and Mesquite cannot, we continued with Bppancestor for further analyses on the full dataset.

Dollo parsimony consistently produced reconstructions of ancestral node gene content that were above the true 5,000 genes threshold (Figure 1A,B). This effect was amplified at nodes closer to the root, where there were more overestimated nodes, and where the estimated gene counts showed the largest inflations above 5,000 (Figure 2, Supplementary Figure 5). The increased overestimates closer to the root that we observed are consistent with our expectations, as nodes closer to the root have more children, and thus more opportunities for non-orthologous sequence homology between pairs of distantly related species to arise, which would then be incorrectly inferred by Dollo parsimony to have been present in their most recent common ancestor. In contrast, maximum likelihood never produced estimated counts above the true 5,000 genes threshold (Figure 1C).

**Figure 1.**
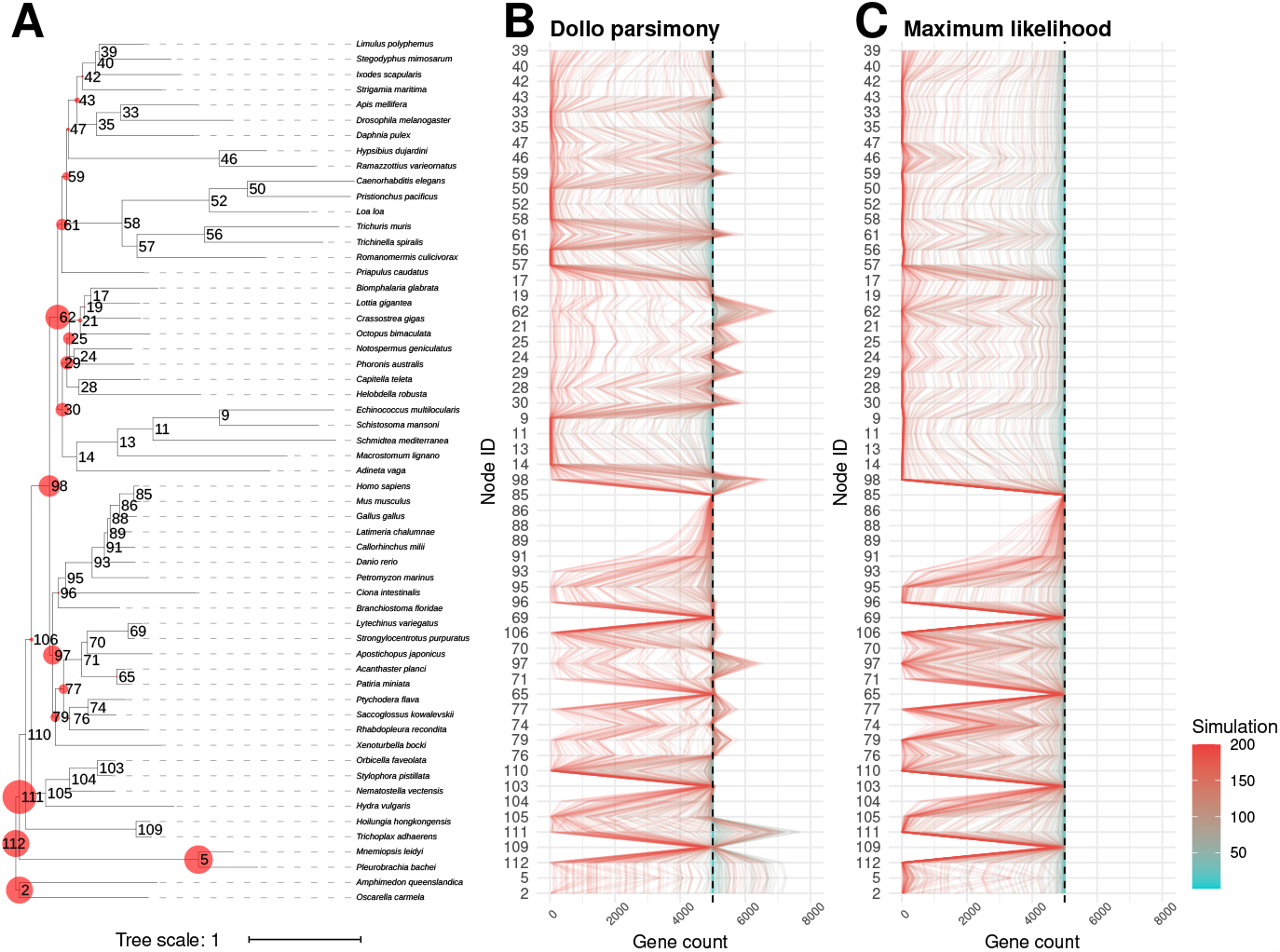
Ancestral gene content reconstructions on a simulated data set containing exactly 5000 orthologs. (A) Phylogenetic tree (Natsidis et al., 2021) depicting the relationship species used in the simulations, highlighting ancestral nodes that were overestimated by Dollo parsimony in at least one case (red circles). The size of the red circles is proportional to the largest number (among all simulations) of estimated genes exceeding 5000. Internal nodes are identified by numbers, which correspond among panels. (B) Gene counts at internal nodes by Dollo parsimony. (C) Gene counts at internal nodes inferred by maximum likelihood. In panels B and C, each line represents the set of inferences from one simulation. Simulation numbers correspond to the rate of sequence evolution used to produce simulated data (lower numbers have lower rates).

**Figure 2.**
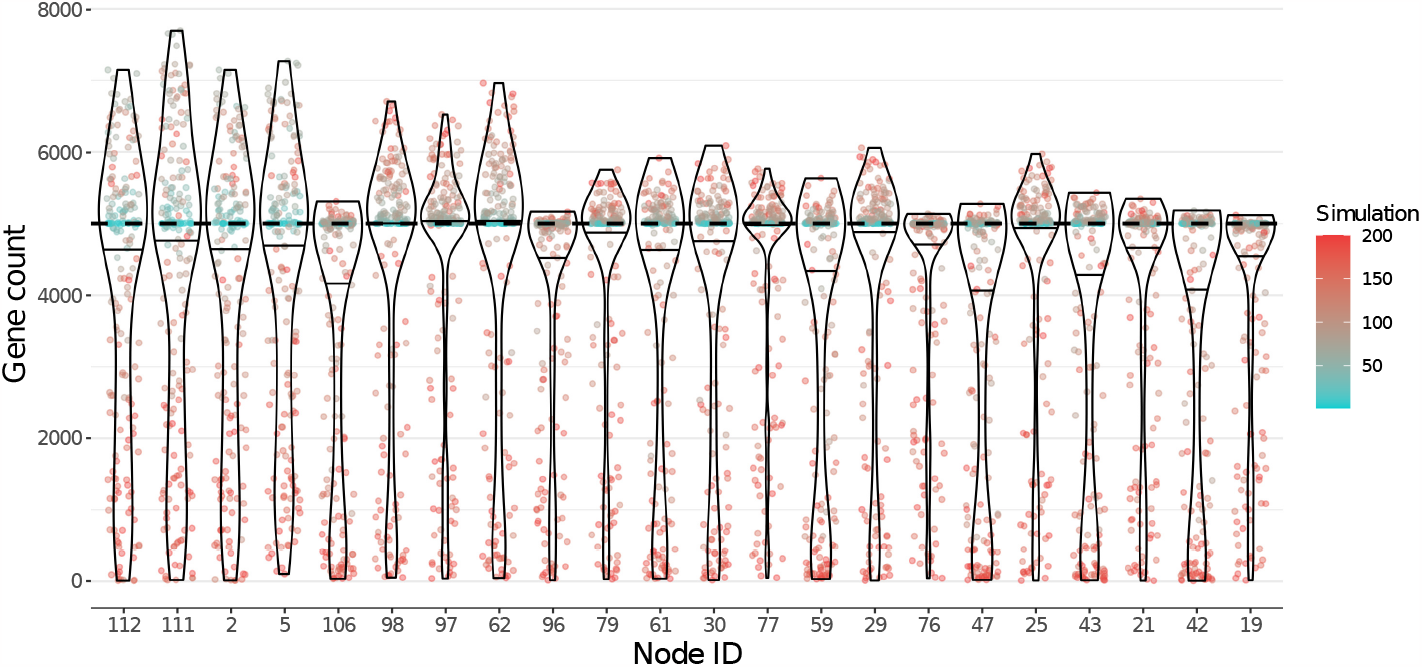
Distributions of inferred gene counts at ancestral nodes that showed the highest levels of overestimation by Dollo parsimony. Each colored dot in a distribution represents the ancestral gene count inferred from one simulation. Simulation numbers correspond to the rate of sequence evolution used to produce simulated data (lower numbers have lower rates). The color scale is identical to that of Figure 1. The nodes on the horizontal axis are ordered by their proximity to the root of the topology (nodes that are closer to the root appear towards the left). The proximity to the root is measured as the number of internal nodes between the node of interest the root and of the tree. This figure shows nodes that were overestimated by at least 105 genes for at least one simulation; distributions for all internal nodes are shown in Supplementary Figure 5.

For Dollo parsimony, both relatively slow and relatively fast evolving simulations produced overestimations (Figure 1B, Figure 2). For maximum likelihood, inferences from slow-evolving simulations were generally close to the true gene count, whereas fast evolving simulations resulted in larger distortions, but never produced any overestimation. The low estimates that Bppancestor produced with fast evolving simulations could be explained by the already underestimated input. Areas of the topology with an accurate input count generated more accurate inferences than areas of the topology with a distorted (underestimated) input count (Supplementary Figure 3).

### Dollo parsimony produces substantially higher estimates of Pfam domain content in the earliest eukaryotes

To contrast the inferences of Dollo parsimony and maximum likelihood on a real dataset, we chose to return to an analysis first carried out in 2011 by Zmasek and Godzik (Zmasek & Godzik, 2011). Their aim was to reconstruct the evolution of Pfam protein domain content in eukaryotes, by sampling the available genomes of a diversity of extant species and applying Dollo parsimony to reconstruct the history of domain gain and loss. We repeated their analysis with an updated set of species from EukProt v3 (Richter et al., 2022) and compared the results of Dollo parsimony versus maximum likelihood on the set of Pfam domains annotated to be present in each species.

Dollo parsimony and maximum likelihood produced substantially different estimates of Pfam domain content at ancestral nodes, as well as counts of domain gain and loss across the eukaryotic tree (Figure 3). Dollo parsimony produced much larger domain counts than maximum likelihood. The estimates from Dollo parsimony also increased in size with proximity to the root, similar to what we observed in our analysis of simulated data. In fact, Dollo parsimony reconstructed a LECA (last eukaryotic common ancestor) with a higher Pfam domain content than any extant eukaryote, which represents almost two times the domain content of the highest estimate from maximum likelihood at any ancestral node. Dollo parsimony displayed a clear tendency towards domain loss versus domain gain (45,723 total losses against 872 total gains). In contrast, the results from maximum likelihood were more balanced between domain gain and domain loss (3,706 total losses against 4,829 total gains) (Figure 3 and Supplementary Figure 6). We also observed a major difference regarding where domain gains occurred in the tree: in Dollo parsimony, most of the domain gains are inferred close to the LECA (as we can see in clades such as Diaphoretickes), whereas maximum likelihood infers most of the domain gains closer to the leaves of the tree (as can be seen in clades such as Amoebozoa) (Supplementary Figure 6).

**Figure 3.**
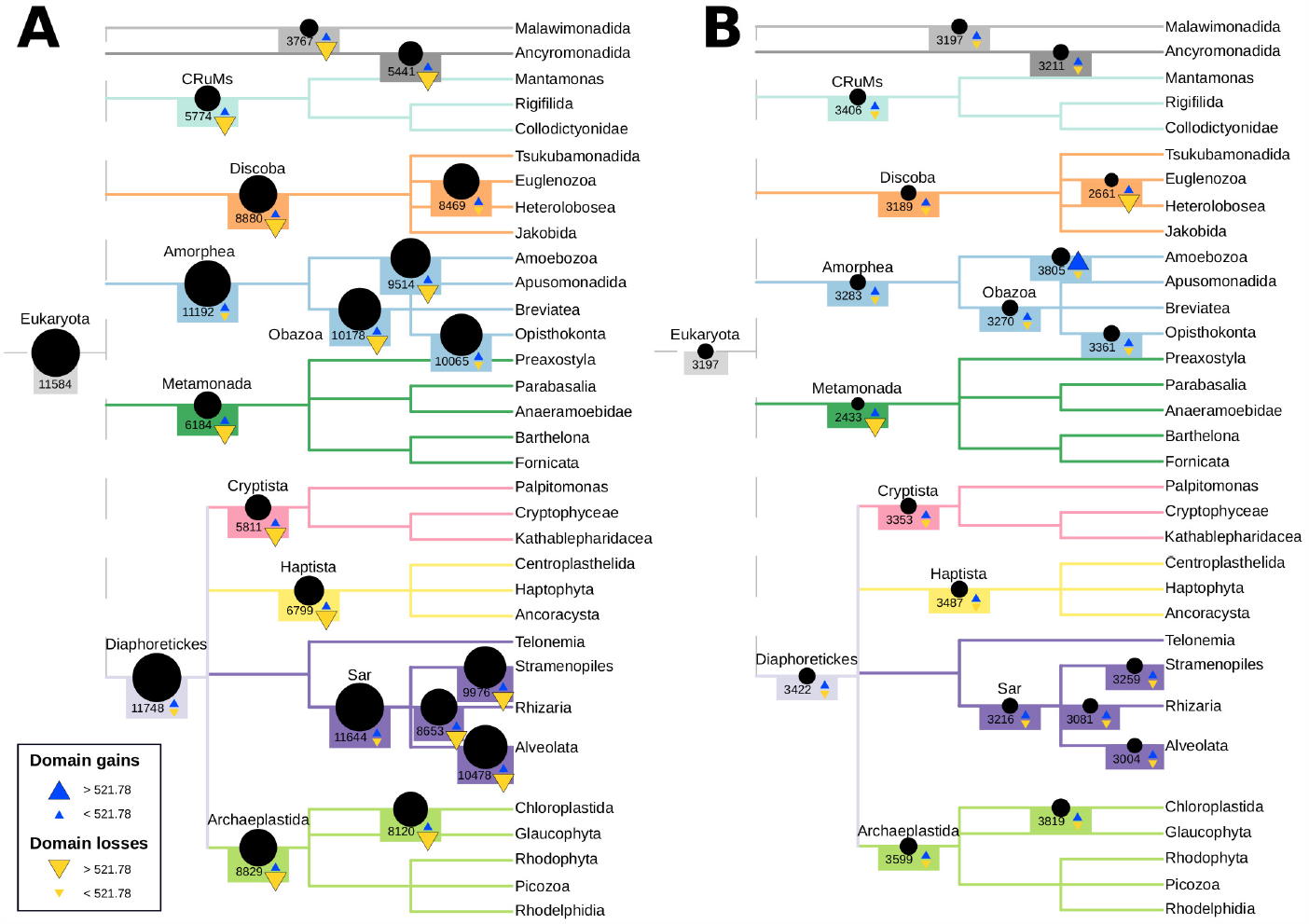
Pfam protein domain counts, gains and losses during eukaryotic evolution, inferred by (A) Dollo parsimony and (B) maximum likelihood. The sizes of the black circles are proportional to the estimated count of domains present at selected nodes. Blue triangles represent inferred protein domain gains, while yellow triangles represent inferred protein domain losses. The threshold separating the two different sizes of triangles is derived from the 3rd quartile of all gain and loss inferences (Q3 = 521.78). Numeric counts of gains and losses are shown in Supplementary Figure 6. Tree topology and node names are derived from UniEuk (Berney et al. 2017),

Our Dollo parsimony inferences are coherent with the results obtained by Zmasek and Godzik. Both studies produced a loss-dominated reconstruction of eukaryotic evolution, with relatively few exceptions. A large protein domain repertoire in the LECA, larger than any extant species, was also inferred in both studies. Even though these two inferences coincide, they are inconsistent with our maximum likelihood reconstruction, which displayed a balance between domain gain and loss, and inferred a LECA with a smaller number of unique Pfam domains than many extant species.

## DISCUSSION

In this study, we tested the hypothesis that Dollo parsimony overestimates ancestral gene content reconstructions, by using simulated data as an input. Next, we analyzed a real data set of Pfam protein domain content across eukaryotes with Dollo parsimony and with maximum likelihood, which did not show evidence for overestimations on simulated data.

The use of simulated data without gene gains or losses, in the first section of this study, provided Dollo parsimony with a “favorable” scenario where no convergent gains, non-orthologous homology (e.g., gene families with ancient duplications) or horizontal gene transfers could generate overestimations in inferred ancestral gene counts. Moreover, the gene counts of extant species provided as input were substantially underestimated for the fastest evolving simulations.

Even in this favorable scenario for Dollo parsimony, we found a clear tendency of Dollo parsimony to overestimate both gene content, as many values were inferred to be higher than the true ancestral value, and gene loss, as the overestimations were larger towards the root of the tree topology. These overestimations result only from orthology inference errors (the splitting of the original orthogroups and random sequence similarity) and could be much more pronounced in a real case, where secondary gains, non-orthologous homology and horizontal gene transfer play a role.

The results of Dollo parsimony were more accurate on slower versus faster-evolving simulations. In faster-evolving simulations, true orthologs were split across multiple orthogroups in Dollo’s input data, resulting in overestimated inferences, as ancestral nodes contained artificially-generated orthogroups in addition to the true ones (Supplementary Figure 2). Beginning with a larger number of input orthogroups in an ancestral reconstruction performed with Dollo parsimony increases the potential number of inferred genes at ancestral nodes. If multiple artificially-generated orthogroups for a single group of true orthologs each contain representative genes from phylogenetically distant species, then inflation of ancestral counts would occur at those species’ last common ancestors. Although this aspect of the input data led to inflation in Dollo parsimony’s ancestral gene content estimates, not all aspects are likely to lead to inflated estimates. In fact, two other aspects would be expected to produce reductions. First, with increasing evolutionary rates, the likelihood also increases that simulated sequences are so distant that they are excluded entirely from any orthogroup (i.e., they are singletons); this would lead to the associated genes being considered to be absent in the species. Second, Dollo parsimony’s input data are binary (either present or absent); therefore, we only accounted for presence or absence of any sequence from a given orthogroup in a species’ proteome, not the number of sequences. When multiple sequences from the same species were incorrectly partitioned into the same orthogroup, they would only be counted once, thereby reducing the total number of input gene presences.

Although Dollo parsimony produced estimates of ancestral gene content that were above the input value of 5,000 in many cases, its inferences in other cases were not inflated above 5,000 and were closer than those of maximum likelihood to the true input value (Supplementary Figure 7). We believe that this is an artifact of the input data to ancestral reconstruction. Dollo parsimony generally produced overestimated inferences across our data set. When presented with underestimated input values, it produced ancestral reconstructions that were overestimates of these underestimates. Therefore, although Dollo parsimony generated clear overestimations, it might have the effect of compensating for underestimations in input data for some cases of ancestral reconstruction.

Our results also have implications for the phylostratigraphy approach, which also relies on Dollo parsimony’s assumption that a gene may only be gained once over evolutionary history, although we note that both gene age overestimation (as might be expected to occur given our results) and gene age underestimation within the phylostratigraphy approach have been extensively debated in the literature (Casola, 2018; Domazet-Lošo et al., 2017; Moyers & Zhang, 2015, 2016, 2017, 2018).

In the second part of our study, we observed strikingly different estimates of ancestral eukaryote Pfam protein domain content when reconstructed with Dollo parsimony versus maximum likelihood. These contradictory results indicate that conclusions of evolutionary studies based on ancestral reconstruction can be extremely dependent on the methodology used.

Our ancestral reconstruction using Dollo parsimony inferred that the last eukaryotic common ancestor had more Pfam domains than any extant eukaryote, and that the evolutionary history of Pfam domains in eukaryotes was dominated by loss. These results are consistent with those of a previous study based on a smaller number of datasets available at the time (Zmasek & Godzik, 2011), and may either reflect the true evolutionary history of Pfam domains in eukaryotes, or may be the result of distortions due to Dollo parsimony. The fact that we demonstrated Dollo parsimony’s inherent tendency to overestimate both ancestral gene content and the number of gene losses using simulated data as input suggests that the evolutionary scenario inferred by Dollo parsimony may have been an artifact of the methodology that was applied.

In the context of our analysis, it is relevant to remark that Pfam profile HMMs are derived from a representative alignment of select taxa, which results in a biased protein domain detection towards biomedically relevant species and an underestimation of detected domains in non-model species (Tassia et al., 2021). Another possible bias in our inference might be caused by the heuristic e-value thresholds implemented in InterProScan (Jones et al., 2014) to assess protein domain presence or absence in input species’ proteomes. These phenomena may help explain the relatively low estimates of domain presence and high numbers of domain losses inferred by both ancestral reconstruction methods in groups that are poorly represented in protein sequence databases (e.g., Metamonada).

Dollo parsimony, and other phylogenetic inference methods and programs involved in the process of ancestral reconstruction, induce biases in the inference of gene content. Therefore, we propose that, in order to mitigate the effects of these biases, the results of different methods should be contrasted in order to assess which ancestral reconstructions are more likely to be an accurate representation of the evolution of the studied organisms rather than an artifact of the methodology. Some alternatives to Dollo parsimony that could be used and compared with each other in ancestral reconstruction studies are programs such as Bppancestor and Mesquite (maximum likelihood) (Guéguen et al., 2013; Maddison, W. P. and D.R. Maddison, 2023), Count (Wagner parsimony and linear birth-death-immigration method) (Csűös, 2010) or MrBayes (Bayesian inference) (Ronquist et al., 2012). Gene tree/species reconciliation methods, such as ALE (Szöllösi et al., 2013; Szöllősi et al., 2015), could also enhance these analyses, as they can also detect horizontal gene transfer events. The orthology inference performed by OrthoFinder2 (Emms & Kelly, 2019) also added some degree of distortion to our input data. Alternative orthology inference methods such as Broccoli (mixed phylogeny-network approach) (Derelle et al., 2020) could be used as valuable comparisons to OrthoFinder2.

Overall, our results indicate that, in ancestral reconstruction studies based on sequence homology, Dollo parsimony tends to overestimate both ancestral gene content and gene loss, consequently, the results of different phylogenetic inference methods should be compared in order to obtain a coherent portrait of evolutionary history.

## DATA AND CODE AVAILABILITY

Configuration files for Bppancestor and Bppml, as well as phylogenetic trees and input files used for ancestral reconstructions are available on GitHub at https://github.com/beaplab/Ancestral-Reconstruction.

## REFERENCES

Altschul, S. F., Gish, W., Miller, W., Myers, E. W., & Lipman, D. J. (1990). Basic local alignment search tool. Journal of Molecular Biology, 215(3), 403–410. 10.1016/S0022-2836(05)80360-2

Berney, C., Ciuprina, A., Bender, S., Brodie, J., Edgcomb, V., Kim, E., Rajan, J., Parfrey, L. W., Adl, S., Audic, S., Bass, D., Caron, D. A., Cochrane, G., Czech, L., Dunthorn, M., Geisen, S., Glöckner, F. O., Mahé, F., Quast, C.,… de Vargas, C. (2017). UniEuk: Time to Speak a Common Language in Protistology! Journal of Eukaryotic Microbiology, 64(3), 407–411. 10.1111/jeu.12414

Bowles, A. M. C., Bechtold, U., & Paps, J. (2020). The Origin of Land Plants Is Rooted in Two Bursts of Genomic Novelty. Current Biology, 30(3), 530-536.e2. 10.1016/j.cub.2019.11.090

Cho, C. H., Park, S. I., Huang, T.-Y., Lee, Y., Ciniglia, C., Yadavalli, H. C., Yang, S. W., Bhattacharya, D., & Yoon, H. S. (2023). Genome-wide signatures of adaptation to extreme environments in red algae. Nature Communications, 14(1), Article 1. 10.1038/s41467-022-35566-x

Csűös, M. (2010). Count: Evolutionary analysis of phylogenetic profiles with parsimony and likelihood. Bioinformatics, 26(15), 1910–1912. 10.1093/bioinformatics/btq315

Csuros, M., Rogozin, I. B., & Koonin, E. V. (2011). A Detailed History of Intron-rich Eukaryotic Ancestors Inferred from a Global Survey of 100 Complete Genomes. PLoS Computational Biology, 7(9), e1002150. 10.1371/journal.pcbi.1002150

Czech, L., Barbera, P., & Stamatakis, A. (2020). Genesis and Gappa: Processing, analyzing and visualizing phylogenetic (placement) data. Bioinformatics, 36(10), 3263–3265. 10.1093/bioinformatics/btaa070

Derelle, R., Philippe, H., & Colbourne, J. K. (2020). Broccoli: Combining Phylogenetic and Network Analyses for Orthology Assignment. Molecular Biology and Evolution, 37(11), 3389–3396. 10.1093/molbev/msaa159

Dollo, L. (1893). Les lois de l’évolution. Bulletin de La Société Belge de Géologie de Paléontologie et d’Hydrologie, 7, 164–166.

Domazet-Loso, T., Brajković, J., & Tautz, D. (2007). A phylostratigraphy approach to uncover the genomic history of major adaptations in metazoan lineages. Trends in Genetics: TIG, 23(11), 533–539. 10.1016/j.tig.2007.08.014

Domazet-Lošo, T., & Tautz, D. (2010). Phylostratigraphic tracking of cancer genes suggests a link to the emergence of multicellularity in metazoa. BMC Biology, 8(1), 66. 10.1186/1741-7007-8-66

Doronina, L., Churakov, G., Kuritzin, A., Shi, J., Baertsch, R., Clawson, H., & Schmitz, J. (2017). Speciation network in Laurasiatheria: Retrophylogenomic signals. Genome Research, 27(6), 997–1003. 10.1101/gr.210948.116

Emms, D. M., & Kelly, S. (2019). OrthoFinder: Phylogenetic orthology inference for comparative genomics. Genome Biology, 20(1), 238. 10.1186/s13059-019-1832-y

Fairclough, S. R., Chen, Z., Kramer, E., Zeng, Q., Young, S., Robertson, H. M., Begovic, E., Richter, D. J., Russ, C., Westbrook, M. J., Manning, G., Lang, B. F., Haas, B., Nusbaum, C., & King, N. (2013). Premetazoan genome evolution and the regulation of cell differentiation in the choanoflagellate Salpingoeca rosetta. Genome Biology, 14(2), R15. 10.1186/gb-2013-14-2-r15

Felsenstein, J. (1983). Parsimony in Systematics: Biological and Statistical Issues. Annual Review of Ecology and Systematics, 14(1), 313–333. 10.1146/annurev.es.14.110183.001525

Groussin, M., Daubin, V., Gouy, M., & Tannier, E. (2016). Ancestral Reconstruction: Theory and Practice. In Encyclopedia of Evolutionary Biology (pp. 70–77). Elsevier. 10.1016/B978-0-12-800049-6.00166-9

Guéguen, L., Gaillard, S., Boussau, B., Gouy, M., Groussin, M., Rochette, N. C., Bigot, T., Fournier, D., Pouyet, F., Cahais, V., Bernard, A., Scornavacca, C., Nabholz, B., Haudry, A., Dachary, L., Galtier, N., Belkhir, K., & Dutheil, J. Y. (2013). Bio++: Efficient Extensible Libraries and Tools for Computational Molecular Evolution. Molecular Biology and Evolution, 30(8), 1745–1750. 10.1093/molbev/mst097

Harms, M. J., & Thornton, J. W. (2010). Analyzing protein structure and function using ancestral gene reconstruction. Current Opinion in Structural Biology, 20(3), 360–366. 10.1016/j.sbi.2010.03.005

Inkscape Project. (2020). Inkscape. Retrieved from https://inkscape.org

Jones, P., Binns, D., Chang, H.-Y., Fraser, M., Li, W., McAnulla, C., McWilliam, H., Maslen, J., Mitchell, A., Nuka, G., Pesseat, S., Quinn, A. F., Sangrador-Vegas, A., Scheremetjew, M., Yong, S.-Y., Lopez, R., & Hunter, S. (2014). InterProScan 5: Genome-scale protein function classification. Bioinformatics, 30(9), 1236–1240. 10.1093/bioinformatics/btu031

Letunic, I., & Bork, P. (2007). Interactive Tree Of Life (iTOL): An online tool for phylogenetic tree display and annotation. Bioinformatics, 23(1), 127–128. 10.1093/bioinformatics/btl529

Li, X., Yang, J.-B., Wang, H., Song, Y., Corlett, R. T., Yao, X., Li, D.-Z., & Yu, W.-B. (2021). Plastid NDH Pseudogenization and Gene Loss in a Recently Derived Lineage from the Largest Hemiparasitic Plant Genus Pedicularis (Orobanchaceae). Plant and Cell Physiology, 62(6), 971–984. 10.1093/pcp/pcab074

Maddison, W. P. and D.R. Maddison. (2023). Mesquite: A modular system for evolutionary analysis. Version 3.81. http://www.mesquiteproject.org

Mans, B. J., de Castro, M. H., Pienaar, R., de Klerk, D., Gaven, P., Genu, S., & Latif, A. A. (2016). Ancestral reconstruction of tick lineages. Ticks and Tick-Borne Diseases, 7(4), 509–535. 10.1016/j.ttbdis.2016.02.002

Najle, S. R., Grau-Bové, X., Elek, A., Navarrete, C., Cianferoni, D., Chiva, C., Cañas-Armenteros, D., Mallabiabarrena, A., Kamm, K., Sabidó, E., Gruber-Vodicka, H., Schierwater, B., Serrano, L., & Sebé-Pedrós, A. (2023). Stepwise emergence of the neuronal gene expression program in early animal evolution. Cell, S0092867423009170. 10.1016/j.cell.2023.08.027

Natsidis, P., Kapli, P., Schiffer, P. H., & Telford, M. J. (2021). Systematic errors in orthology inference and their effects on evolutionary analyses. iScience, 24(2), 102110. 10.1016/j.isci.2021.102110

Paps, J., & Holland, P. W. H. (2018). Reconstruction of the ancestral metazoan genome reveals an increase in genomic novelty. Nature Communications, 9(1), Article 1. 10.1038/s41467-018-04136-5

Park, H., Park, J. H., Lee, Y., Woo, D. U., Jeon, H. H., Sung, Y. W., Shim, S., Kim, S. H., Lee, K. O., Kim, J.-Y., Kim, C.-K., Bhattacharya, D., Yoon, H. S., & Kang, Y. J. (2021). Genome of the world’s smallest flowering plant, Wolffia australiana, helps explain its specialized physiology and unique morphology. Communications Biology, 4(1), Article 1. 10.1038/s42003-021-02422-5

Pérez, F., Arroyo, M. T. K., Medel, R., & Hershkovitz, M. A. (2006). Ancestral reconstruction of flower morphology and pollination systems in Schizanthus (Solanaceae). American Journal of Botany, 93(7), 1029–1038. 10.3732/ajb.93.7.1029

R Core Team. (2022). R: A language and environment for statistical computing. R Foundation for Statistical Computing, Vienna, Austria.

Repetti, S. I., Iha, C., Uthanumallian, K., Jackson, C. J., Chen, Y., Chan, C. X., & Verbruggen, H. (2022). Nuclear genome of a pedinophyte pinpoints genomic innovation and streamlining in the green algae. New Phytologist, 233(5), 2144–2154. 10.1111/nph.17926

Richter, D. J., Berney, C., Strassert, J. F. H., Poh, Y.-P., Herman, E. K., Muñoz-Gómez, S. A., Wideman, J. G., Burki, F., & de Vargas, C. (2022). EukProt: A database of genome-scale predicted proteins across the diversity of eukaryotes. Peer Community Journal, 2. 10.24072/pcjournal.173

Ronquist, F., Teslenko, M., van der Mark, P., Ayres, D. L., Darling, A., Höhna, S., Larget, B., Liu, L., Suchard, M. A., & Huelsenbeck, J. P. (2012). MrBayes 3.2: Efficient Bayesian Phylogenetic Inference and Model Choice Across a Large Model Space. Systematic Biology, 61(3), 539–542. 10.1093/sysbio/sys029

Ros-Rocher, N., Pérez-Posada, A., Leger, M. M., & Ruiz-Trillo, I. (2021). The origin of animals: An ancestral reconstruction of the unicellular-to-multicellular transition. Open Biology, 11(2), 200359. 10.1098/rsob.200359

RStudio Team. (2020). RStudio: Integrated Development for R. RStudio, PBC, Boston, MA. http://www.rstudio.com/

Stevenson, K., Hughes, V. M., de Juan, L., Inglis, N. F., Wright, F., & Sharp, J. M. (2002). Molecular Characterization of Pigmented and Nonpigmented Isolates of Mycobacterium avium subsp. Paratuberculosis. Journal of Clinical Microbiology, 40(5), 1798–1804. 10.1128/JCM.40.5.1798-1804.2002

Sverdlov, A. V., Rogozin, I. B., Babenko, V. N., & Koonin, E. V. (2004). Reconstruction of Ancestral Protosplice Sites. Current Biology, 14(16), 1505–1508. 10.1016/j.cub.2004.08.027

Szöllősi, G. J., Davín, A. A., Tannier, E., Daubin, V., & Boussau, B. (2015). Genome-scale phylogenetic analysis finds extensive gene transfer among fungi. Philosophical Transactions of the Royal Society B: Biological Sciences, 370(1678), 20140335. 10.1098/rstb.2014.0335

Szöllösi, G. J., Tannier, E., Lartillot, N., & Daubin, V. (2013). Lateral Gene Transfer from the Dead. Systematic Biology, 62(3), 386–397. 10.1093/sysbio/syt003

Tassia, M. G., David, K. T., Townsend, J. P., & Halanych, K. M. (2021). TIAMMAt: Leveraging Biodiversity to Revise Protein Domain Models, Evidence from Innate Immunity. Molecular Biology and Evolution, 38(12), 5806–5818. 10.1093/molbev/msab258

Wall, L., Christiansen, T., & Orwant, J. (2000). Programming Perl.” O’Reilly Media, Inc.”.

Zhou, L., Kappel, F., Wiersma, P. A., Hampson, C., & Bakkeren, G. (2005). GENETIC ANALYSIS AND DNA FINGERPRINTING OF SWEET CHERRY CULTIVARS AND SELECTIONS USING AMPLIFIED FRAGMENT LENGTH POLYMORPHISMS (AFLP). Acta Horticulturae, 667, 37–44. 10.17660/ActaHortic.2005.667.2

Zhu, Q. (2014). AfterPhylo. A Perl script for manipulating trees after phylogenetic reconstruction. https://github.com/qiyunzhu/AfterPhylo/. 0.9.1 ed

Zmasek, C. M., & Godzik, A. (2011). Strong functional patterns in the evolution of eukaryotic genomes revealed by the reconstruction of ancestral protein domain repertoires. Genome Biology, 12(1), R4. 10.1186/gb-2011-12-1-r4

