## Supplementary Figures for "Dollo parsimony overestimates ancestral gene content reconstructions"

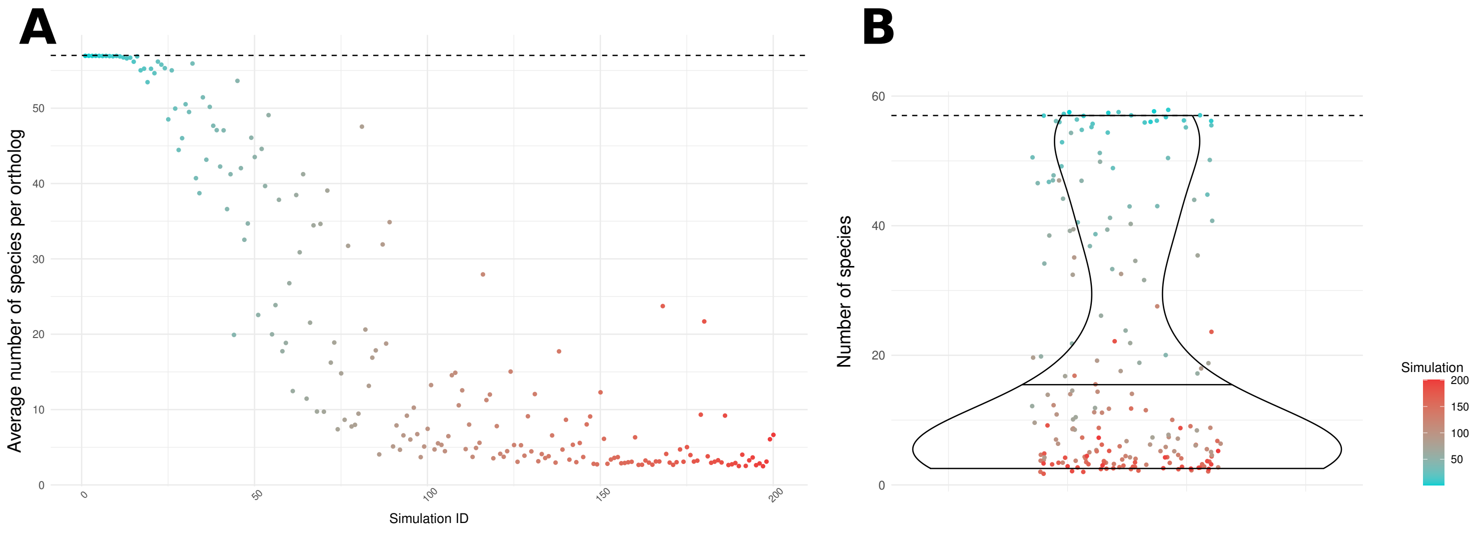


**Supplementary Figure 1. Number of species per orthogroup used as input to ancestral gene content reconstruction.** (A) Average number of species per orthogroup per simulation. (B) Distribution of the average number of species per orthogroup per simulation. In both panels, simulation numbers correspond to the rate of sequence evolution used to produce simulated data (lower numbers have lower rates). The dashed line represents the correct simulated value of 57 species.

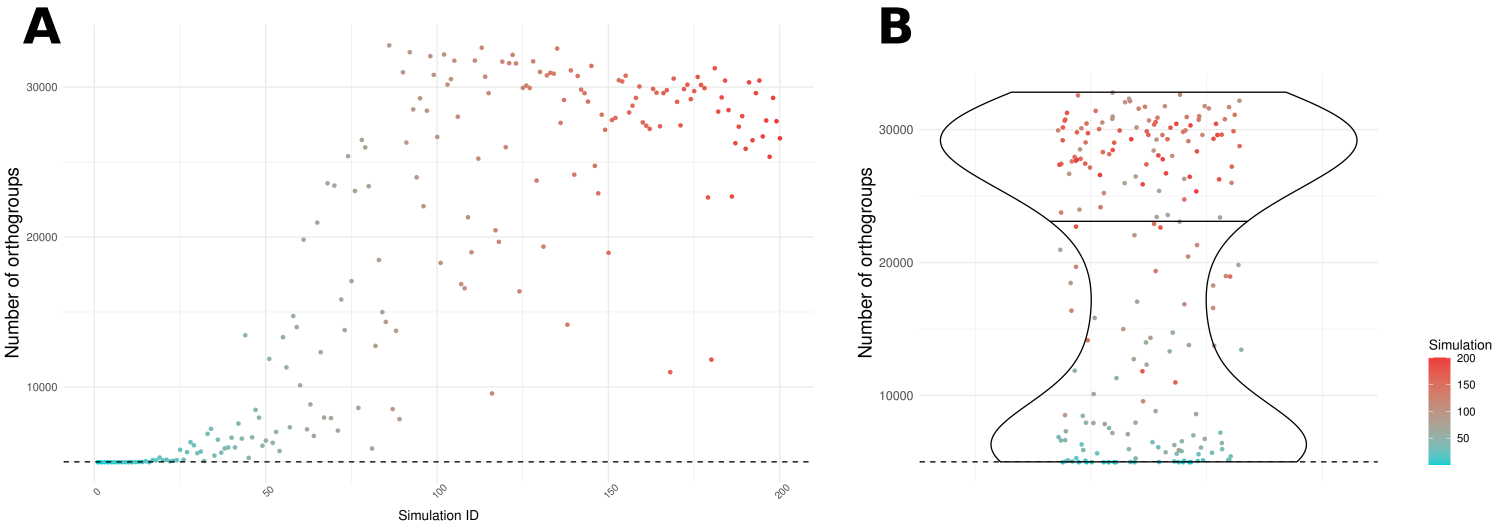


**Supplementary Figure 2. Input orthogroups to ancestral gene content reconstruction.** (A) Total number of input orthogroups to ancestral gene content reconstruction by simulation. (B) Distribution of the number of input orthogroups by simulation. In both panels, simulation numbers correspond to the rate of sequence evolution used to produce simulated data (lower numbers have lower rates). The red dashed line represents the correct simulated value of 5000 orthogroups.

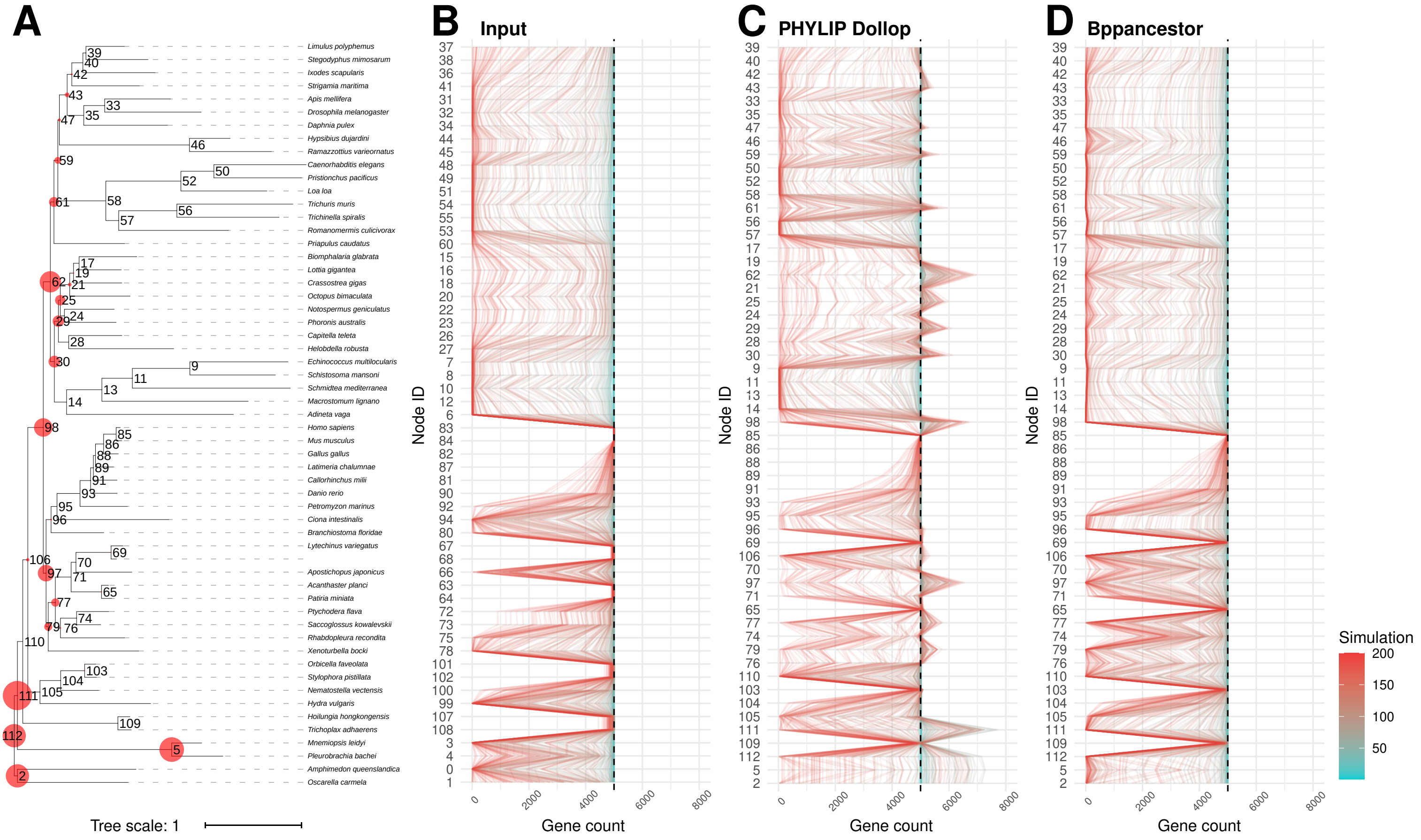


**Supplementary Figure 3. Input and gene count of ancestral gene content reconstructions on a simulated data set containing exactly 5000 orthologs.** (A) Phylogenetic tree (Natsidis et al., 2021) depicting the relationship among species used in the simulations, highlighting ancestral nodes that were overestimated by Dollo parsimony in at least one case (red circles). The size of the red circles is proportional to the largest number (among all simulations) of estimated genes exceeding 5000. Internal nodes are identified by numbers, which correspond among panels. (B) Input counts at all leaf nodes (extant species) for the ancestral reconstruction. (C) Gene count at all internal nodes inferred by Dollo parsimony. (D) Gene count at all internal nodes inferred by Maximum likelihood). For A, each line represents the set of input orthologous groups from one simulation. For B and C each line represents the set of output ancestral gene content inferences from one simulation. Simulation numbers correspond to the rate of sequence evolution used to produce simulated data (lower numbers have lower rates).

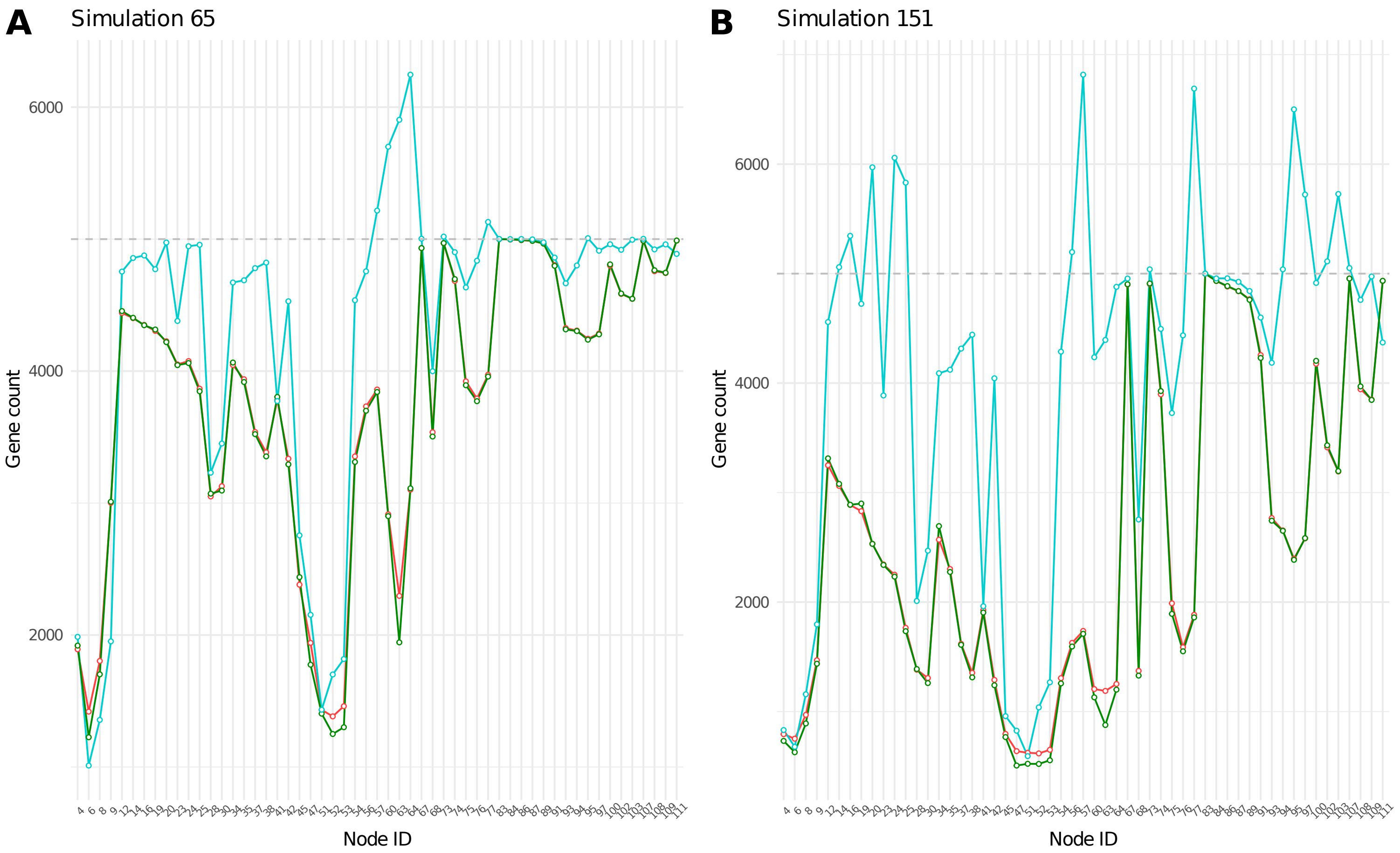


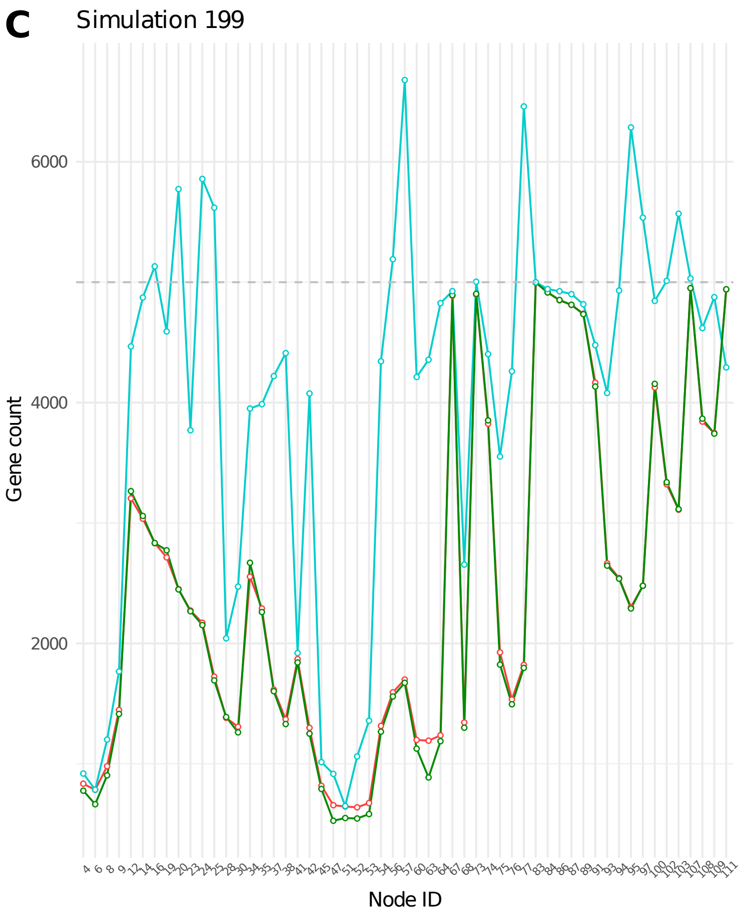


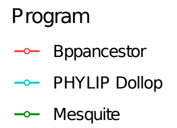


**Supplementary Figure 4. Comparison of two maximum likelihood implementations on three selected simulations.** Gene counts are shown at internal nodes (see Figure 1A) as inferred by Bppancestor and Mesquite, two different maximum likelihood implementations. Results from Dollo parsimony are shown for reference. (A) Simulation 65, (B) Simulation 151 and (C) Simulation 199. Simulation numbers correspond to the rate of sequence evolution used to produce simulated data (lower numbers have lower rates). The gray dashed line indicates the 5000 threshold.

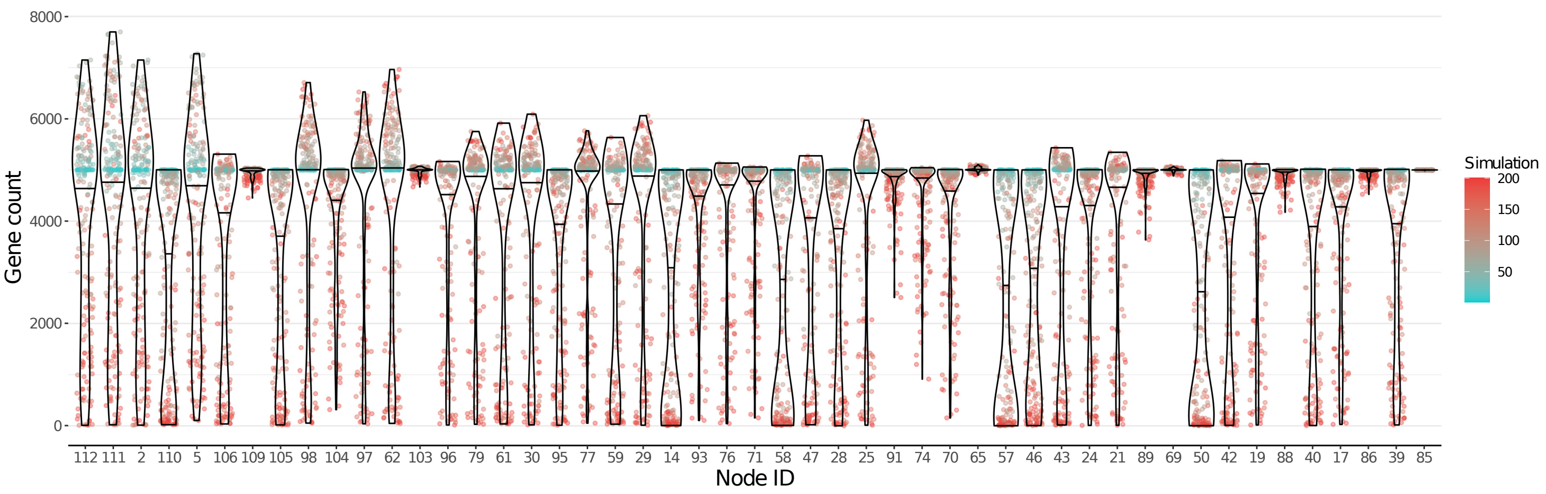


**Supplementary Figure 5. Distribution of the gene count at all ancestral nodes that were overestimated by Dollo parsimony.** Each colored dot in a distribution represents the ancestral gene count inferred from one simulation. Simulation numbers correspond to the rate of sequence evolution used to produce simulated data (lower numbers have lower rates). The nodes in the horizontal axis are ordered by their proximity to the root of the topology, so that nodes that are closer to the root appear towards the left. The proximity to the root is measured as the number of internal nodes between the node of interest and the root of the tree.

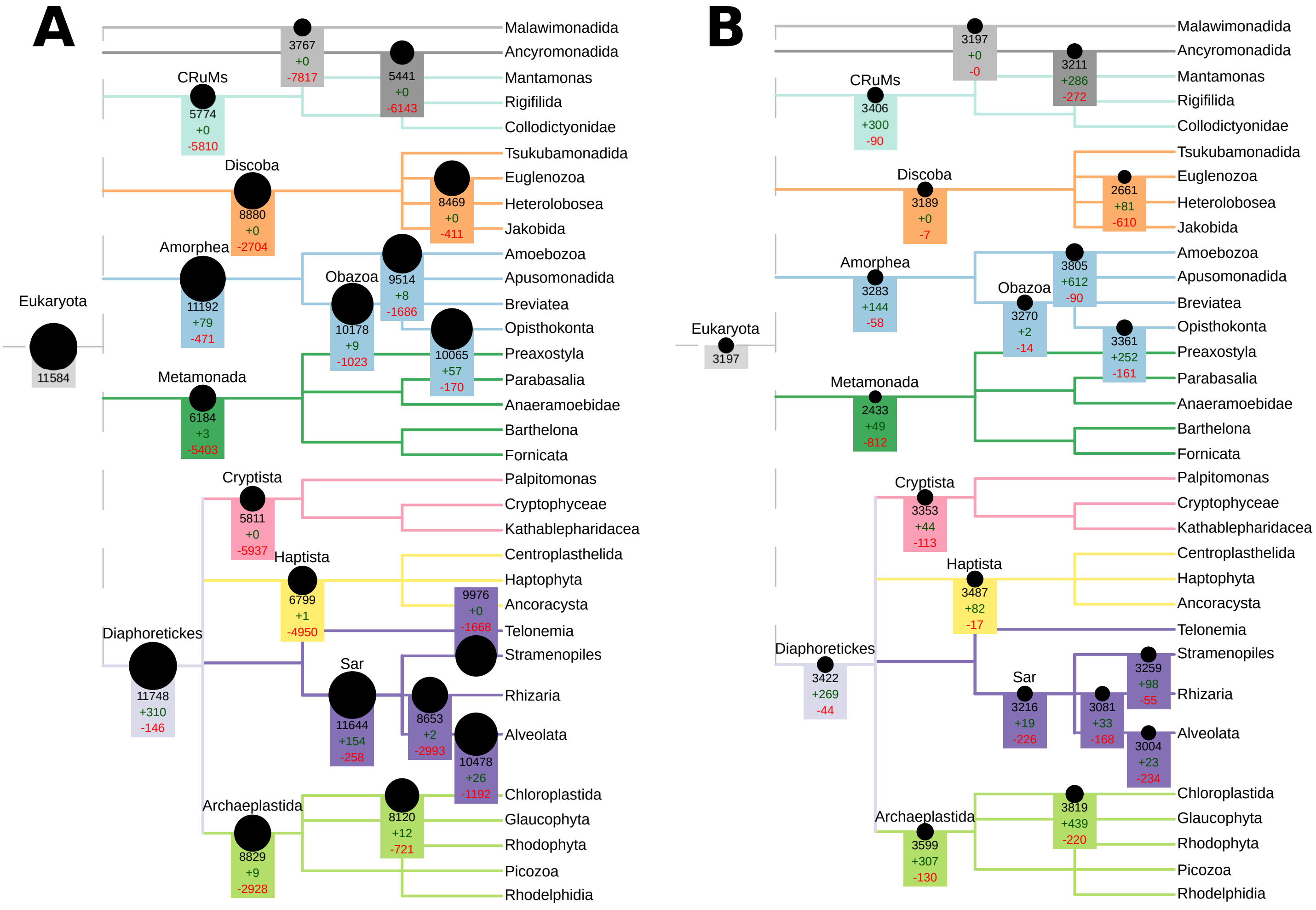


**Supplementary Figure 6. Pfam protein domain counts, gains and losses during eukaryotic evolution, inferred by (A) Dollo parsimony and (B) maximum likelihood.** The sizes of the black circles are proportional to the estimated count of domains present at each node. The number of protein domains present at selected nodes are shown in black, gains are shown in green, and losses are shown in red. Tree topology and node names are derived from UniEuk (Berney et al., 2017).

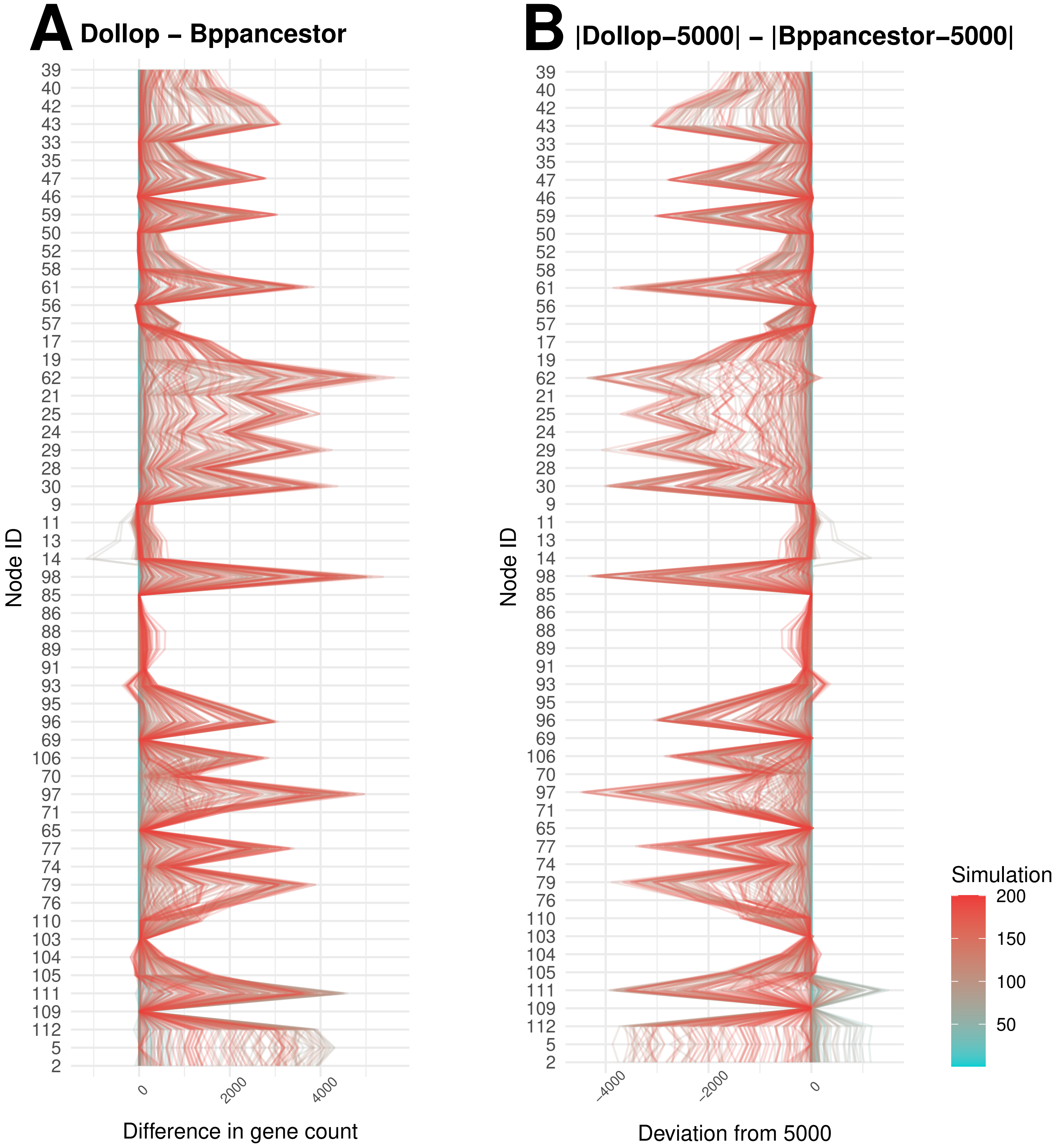


**Supplementary Figure 7.** Direct comparison of Dollo parsimony versus maximum likelihood inferences by simulation. (A) Difference in the inference of Dollo parsimony (Dollop) and maximum likelihood(Bppancestor), expressed as the ancestral gene content at a given node for a given simulation estimated by Dollo parsimony minus that of maximum likelihood. Negative values represent a higher inference from maximum likelihood, while positive values represent a higher inference from Dollo parsimony. (B) Difference in the distances of each program's inference to 5,000 (the true count). Negative values represent a smaller distance from Dollo parsimony's inference to 5,000, while positive values represent a smaller distance from Maximum likelihood's inference to 5,000. For A and B each line represents the set of output ancestral gene content inferences from one simulation. Simulation numbers correspond to the rate of sequence evolution used to produce simulated data (lower numbers have lower rates).
